# caliPER: A software for blood-free parametri*c* P*a*t*l*ak mapp*i*ng using *PE*T/M*R*I input function

**DOI:** 10.1101/2021.07.08.451713

**Authors:** Praveen Dassanayake, Lumeng Cui, Elizabeth Finger, Matthew Kewin, Jennifer Hadaway, Andrea Soddu, Bjoern Jakoby, Sven Zuehlsdorf, Keith S St Lawrence, Gerald Moran, Udunna C Anazodo

## Abstract

Routine clinical use of absolute PET quantification techniques is limited by the need for serial arterial blood sampling for input function and more importantly by the lack of automated pharmacokinetic analysis tools that can be readily implemented in clinic with minimal effort. PET/MRI provides the ability for absolute quantification of PET probes without the need for serial arterial blood sampling using image-derived input functions (IDIFs). Here we introduce caliPER, a modular and scalable software for simplified pharmacokinetic modelling of PET probes with irreversible uptake or binding based on PET/MR IDIFs and Patlak Plot analysis. caliPER generates regional values or parametric maps of net influx rate (*K*_i_) using reconstructed dynamic PET images and anatomical MRI aligned to PET for IDIF vessel delineation. We evaluated the performance of caliPER for blood-free region-based and pixel-wise Patlak analyses of [^18^F] FDG by comparing caliPER IDIF to serial arterial blood input functions and its application in imaging brain glucose hypometabolism in Frontotemporal dementia. IDIFs corrected for partial volume errors including spill-out and spill-in effects were similar to arterial blood input functions with a general bias of around 6-8%, even for arteries <5 mm. The *K*_*i*_ and cerebral metabolic rate of glucose estimated using caliPER IDIF were similar to estimates using arterial blood sampling (<2%) and within limits of whole brain values reported in literature. Overall, caliPER is a promising tool for irreversible PET tracer quantification and can simplify the ability to perform parametric analysis in clinical settings without the need for blood sampling.

**Highlights:**

- caliPER is an adaptable image processing software for extracting image-derived input functions and generating parametric maps of irreversible PET tracer uptake.
- Anatomical (T1-weighted/T2-weighted/time-of-flight angiography) MRI carefully aligned to PET provides a robust approach for delineation of vessels on PET, eliminating the need for serial blood sampling for input functions.
- Application of caliPER in modelling glucose uptake in patients with Frontotemporal dementia, demonstrates the feasibility of absolute quantification of cerebral metabolic rate of glucose in clinical populations.

## 1. Introduction

The superior sensitivity of positron emission tomography (PET) provides a unique approach for probing biological processes in live humans including molecular and neurochemical changes related to disease. The advantage of PET lies in the ability to accurately quantify the concentration of tracers in tissue by modelling the activity in arterial blood and tissue over time, as the tracer is delievered and taken up by tissue. Traditionally, tracer distribution is characterized by semi-quantitative standardized uptake values (SUVs) obtained at a specific time when the tracer concentration is assumed to be in steady state. Because, derivation of SUVs does not require invasive arterial blood sampling and are time efficient with respect to PET image acquisition, SUVs has become the de facto standard for clinical PET quantification. However, SUVs inherently omit specific physiology and biochemical parameters involved with tracer influx and efflux from tissues, limiting PET’s ability to detect subtle changes in tissue -a promising early indicator of disease (Lammertsma, 2017; Wang et al., 2020). And, specifically for application in neurodegenerative disease, the use of SUVs often requires a valid reference region in order to sensitively quantify changes in brain tissue, which can be unavailable for some tracers (e.g. translocator proteins or synaptic vesicle protein 2A PET probes) (McCluskey et al., 2020) and less reliable in some conditions (longitudinal studies or severe cases of dementia) (Mosconi, 2013).

To overcome these drawbacks and exploit the full potential of PET for molecular imaging, several approaches have been proposed to negate the use of invasive serial arterial blood sampling for measuring the arterial input function (AIF) that is required for PET quantification. These include the use of a population-based arterial input function (PBAIF) averaged from normalized AIF of a group of individuals (Takikawa et al., 1993). However, PBAIF are not subject-specific and can introduce errors to quantification due to differences in tracer administration/injection protocols, which usually requires a few blood samples to calibrate or scale to an individual (Croteau et al., 2010). Alternatively, image-derived input functions (IDIF) extracted from the activity within blood vessels captured on dynamic PET images can provide a subject-specific approach for generating input functions without the requirement of blood sampling (blood-free) (Croteau et al., 2010; Zanotti-Fregonara et al., 2009). In order to improve the accuracy of IDIF for PET quantification, careful delineation of arterial vessels on PET images is imperative. The limited spatial resolution of PET can blur measured activity within blood vessels, mixing the signals from the blood with surrounding tissue, especially if the vessel size is comparable with the spatial resolution of PET systems, for example the internal carotid arteries in adults (average diameter ∼5mm) (Krejza et al., 2006). With the increased availability of integrated PET and magnetic resonance imaging (MRI) systems, there is renewed interest in mapping PET’s multi-parametric information, where the higher spatial resolution of simultaneously acquired anatomical MRI can facilitate precise delineation of blood vessels on PET, increasing the accuracy of IDIF (Ringheim et al., 2020; Sari et al., 2016; Sundar et al., 2018). However, to enable routine use of IDIF for absolute parametric mapping of tracer concentration using PET/MRI, simplified approaches for tracer kinetic modelling must be readily available (Wang et al., 2020). Recently, we developed a simplified IDIF approach that combines high-resolution anatomical MRI with dynamic PET data (Anazodo et al., 2015), and demonstrated its feasibility for parametric mapping of [^68^Ga] PSMA-11for prostate cancer imaging (Ringheim et al., 2020).

Here, we present a software prototype for application of this simplified MRI-guided blood-free IDIF approach in region-based and pixel-wise parametric mapping of irreversible PET tracers (e.g. [^18^F]fluorodeoxyglucose (FDG), [^68^Ga]-PSMA-11 and [^18^F]-NaF). This semi-automated prototype, caliPER (parametri**c** P**a**t**l**ak mapp**i**ng using **PE**T/M**R**I input function) 1) uses any anatomical MR image, as a guide to define arterial blood vessels via a region growing segmentation algorithm and a skeletonized multimodal vessel mask registration approach, 2) corrects PET signal contamination (mixture of signal from vessels and surrounding tissue -partial volume errors (PVE) including tracer influx to the vessel from the surrounding tissues -spill-in errors) using simulated correction factors optimized for PET/MR (Croteau et al., 2010; Feng et al., 2012), to enable improved extraction of IDIF from dynamic PET images, and 3) automatically computes parametric PET images using the Patlak graphical analysis method to simplify the complexity of PET compartmental kinetic modelling (Patlak et al., 1983). First, we evaluated the accuracy of our optimized PVE correction method by comparing IDIF extracted from the relatively small brain vessels of a porcine model, to the gold standard AIF. Then, we investigated the performance of the PET/MRI IDIF extracted using caliPER in a cohort of healthy volunteers by comparing the PET/MRI IDIF to a PBAIF (Sundar et al., 2018). Finally, we demonstrate the feasibility of applying caliPER in quantifying the regional cerebral metabolic rate of glucose (CMRglc) in a cohort of patients with frontotemporal dementia (FTD).

## 2. Software Framework

### 2.1 caliPER Overview

caliPER is an investigational adaptable PET/MR image processing prototype tool for extracting MRI-guided IDIF and performing subsequent Patlak graphical analysis to generate region-specific values or pixel-wise parametric maps describing the net uptake rate (*K*_i_) and the distribution volume of unmetabolized tracer in tissue. This prototype was developed on the *syngo*.Via Frontier platform (Siemens Healthineers, Erlangen, Germany) powered by MeVisLab 2.7.1 (MeVis Medical Solutions AG, Bremen, Germany) for fast image processing on multiple operating systems (Microsoft Windows, Linux, or macOS), using an intuitive graphical user interface and a modular scalable network design. caliPER has been tested for analysis of [^68^Ga]-PSMA-11 for primary prostate cancer imaging (Ringheim et al., 2020), [^18^F]-NaF in femoroacetabular impingement syndrome (Cui *et al* 2019), and for FDG imaging of neurodegeneration, as illustrated below.

On the backend, caliPER’s processing pipeline summarized in figure 1, executes a series of steps described below, to output IDIF tabulated data files and parametric maps in DICOM format. The frontend features a user-friendly control panel and interactive display window as illustrated in figure 2 and supplementary figure S1. The control panel provides parameter entry options for optimization of modules in the pipeline e.g. intensity threshold controls for segmentation, blood glucose value, FDG lump constant, etc. The display window accompanies each processing step to offer immediate user feedback and outcomes, with orthogonal viewing or high-quality rendering display options.

**Figure 1:**
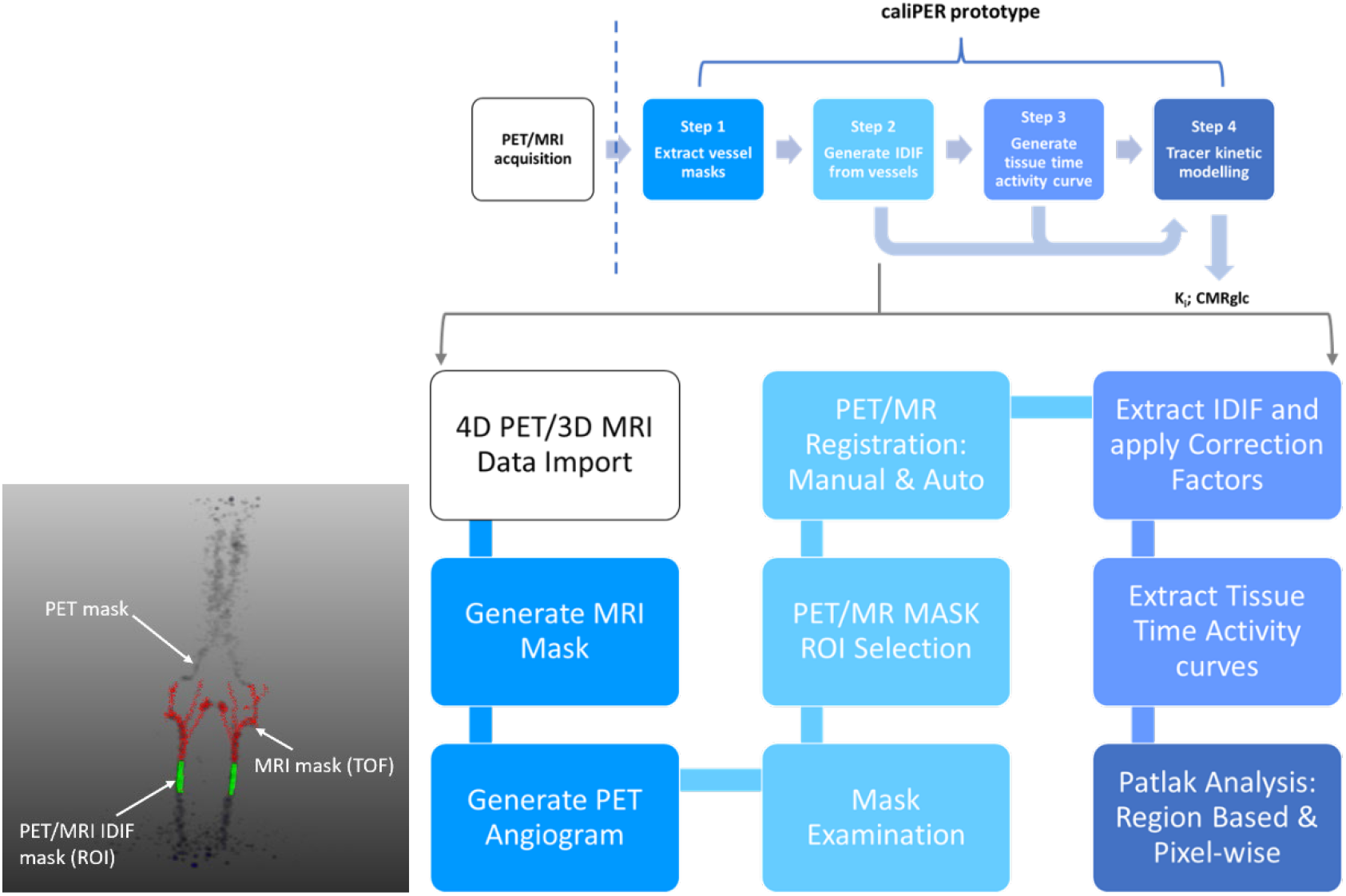
caliPER image processing workflow and functionalities for generating blood-free IDIF and parametric images from reconstructed PET and MRI. An example of co-registered MRI vessel mask of a pig is shown to the left, after MERIT registration and identified PET/MRI mask for automated IDIF extraction.

**Figure 2:**
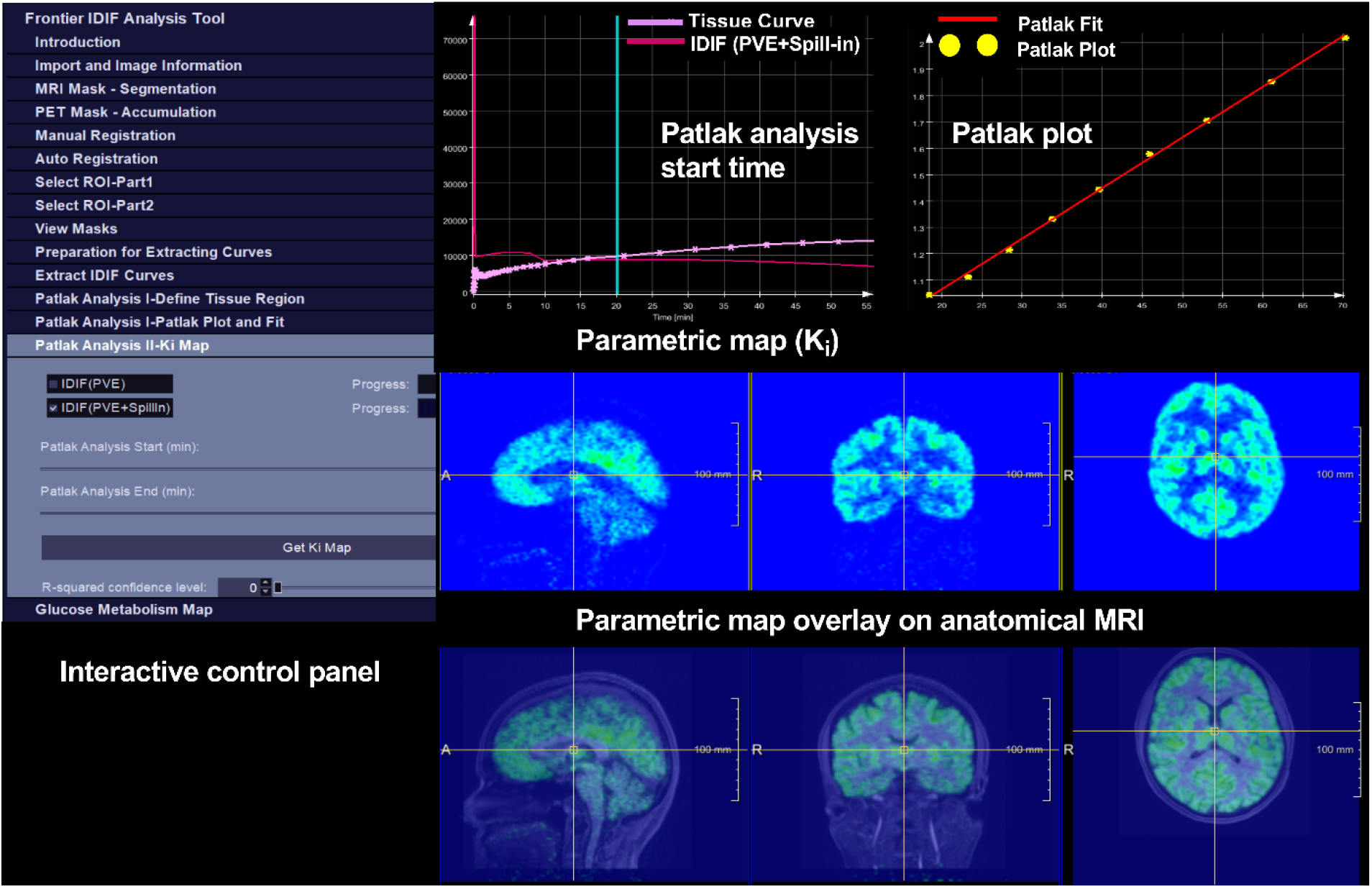
A screen capture of caliPER’s Front-end showcasing one processing step, the user-friendly control panel, and interactive display window. caliPER is an investigational software prototype for analysis of neurological and body PET imaging data.

caliPER requires a dynamic PET time series and an anatomical MRI in DICOM format as inputs for IDIF extraction and parametric mapping. For FDG, an individual’s pre-injection blood glucose can be entered in the control panel to generate CMRglc maps as demonstrated below. In general, caliPER is designed to 1) automatically extracts IDIFs from PET timeseries within vessel masks segmented from anatomical MRI after rigid registration of MRI to PET, 2) correct IDIFs for PVE and errors (Croteau et al., 2010; Feng et al., 2012), and 3) preform region-based or pixel-wise Patlak analysis. These steps are briefly described below.

### 2.2 MRI Arterial Vessel Mask

The MRI vessel mask is generated by segmenting arterial blood vessels on two/three dimensional (2D/3D) anatomical MRI using a seeded region growing segmentation algorithm implemented in MeVisLab. At least one voxel point (seed) within a vessel in a cross-section of the MRI is required to initiate region growing - a search of all contiguous voxels with intensity values that fall within a specified threshold interval and within the immediate vicinity or neighbourhood (x,y,z plane) of the seed. The region growing algorithm segments the vessel tree from background tissue within the specified region and over single or multiple slices. To improve the performance of the region growing segmentation, an image enhancement approach based on a multi-scale vesselness measure (Hessian matrix function) is applied to the MR image, as a pre-processing step before segmentation to enhance the ability to detect vessel boundaries and their tubular-like structures (Friman et al., 2010). An option for manual segmentation with free hand contouring of vessels is included to generate MRI vessel masks if region-growing fails, for example when high resolution anatomical MRI is not available or when arterial vessels cannot be clearly delineated from background tissue. In caliPER, the user initiates region growing simply by placing a seed on the vessel of interest on one or more slice(s) and orientation(s) of the anatomical MRI image.

### 2.3 PET and MR Image Co-registration

The caliPER prototype adopts a two-step registration process, an initial manual registration followed by an automatic registration as shown in figure 1. To improve the accuracy and efficiency of the alignment between the MRI vessel mask and the dynamic PET image volumes, the MRI vessel mask is first manually registered to a PET vessel mask generated by time-averaging PET images over the initial 30 seconds using 3D rigid body registration including translation and rotation. This step is meant to significantly decrease misalignment between the two modalities, providing an accurate initialization of the auto-registration step. Auto-registration and affine-linear transformation of the MRI vessel mask to the PET image space is performed using the **MEVIS Image Registration Toolkit (**MERIT) (Boehler et al., 2011) provided in the MeVisLab developer environment (Heckel et al., 2009). This automated step employs a Newton-type optimizer to further refine the registration by iteratively increasing the similarity between the reference image (PET vessel mask), and the template image (MRI vessel mask). Because, the PET and the MRI vessel mask intensities appear quite similar (near angiograms), the intensity-based sum of squared differences (SSD) was chosen as the most suitable similarity measure.

### 2.4 PET/MR Image-derived Input Functions

The IDIFs are automatically generated by extracting the median PET activity within the co-registered MRI vessel mask at each timeframe. The IDIFs are corrected for PVE both spill-out effects of PET activity within arteries over the surrounding tissue and spill-in effects of PET activity from surrounding tissue into the arteries and for using equation 1 (Chen et al., 1998):

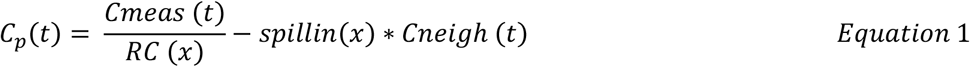

where at any given time *t, Cp* is the ‘true’ tracer concentration in blood (corrected IDIF), *Cmeas* is the measured tracer concentration within the vessel (uncorrected IDIF), *Cneigh* is the tracer concentration in the neighbouring tissues around the vessel, and *RC* and *spillin* are the time-independent recovery coefficient for spill-out and spill-in coefficient respectively at a given vessel diameter (*x)*. The *Cneigh* mask is the difference between the MRI vessel mask dilated by a kernel size of 10 voxel and the non-dilated (i.e., IDIF) vessel mask.

The recovery and spill-in coefficients were estimated as described in (Croteau et al., 2010) and (Feng et al., 2012) using a NEMA IEC Body Phantom Set™ containing a hollow torso that simulates background concentration in human body and refitted six 2.5 cm long cylinders with inner diameters ranging from 5 to 30 mm, designed to mimic the arterial geometry and capture the resolution limit of the internal carotid arteries (supplementary figure S2). A detailed description of the phantom experiments is outlined in the supplementary section. Because the PVE corrections are dependent on the geometry (particularly the size) of the vessel and automatically applied based on the vessel diameter, the vessel masks are restricted to a segment of the vascular tree where the vessels are relatively straight and have the largest possible uniform diameter (figure 1 – ROI selection, PET/MR IDIF mask).

In caliPER, the diameter of the selected PET/MRI IDIF ROI is automatically calculated from the average area of the left and right ROI vessel (supplementary figure S3), or alternatively from one side (left/right) of the vessel in cases where vessel patency is compromised (e.g., stenosis). A two-segment three-gamma fitting model which separately fits the peak and steady-state tail end, can be used to denoise the corrected IDIFs.

### 2.5 Patlak Graphical Analysis Implementation

A fully automated pixel-wise and region-based parametric analysis can be performed within caliPER for tracers with irreversible binding/uptake. This is based on implementation of the Patlak graphical analysis approach for simplified tracer kinetic modelling of tracers with irreversible binding/uptake (Patlak et al., 1983).

The Patlak graphical analysis approach describes the linear transfer kinetics of a two-tissue compartment model (figure 3) consisting of a blood plasma compartment (*C*_*p*_), a non-specifically bound or free tissue compartment (*C*_*1*_), and a trapped or irreversible tissue compartment (*C*_*2*_) with the assumption that the reversible tissue compartment is in equilibrium with the blood plasma compartment after sufficient time post-injection. Based on this assumption, the net uptake rate or total volume of distribution of the PET tracer in the tissue of interest can be estimated using equation 2 (Patlak et al., 1983):

**Figure 3:**
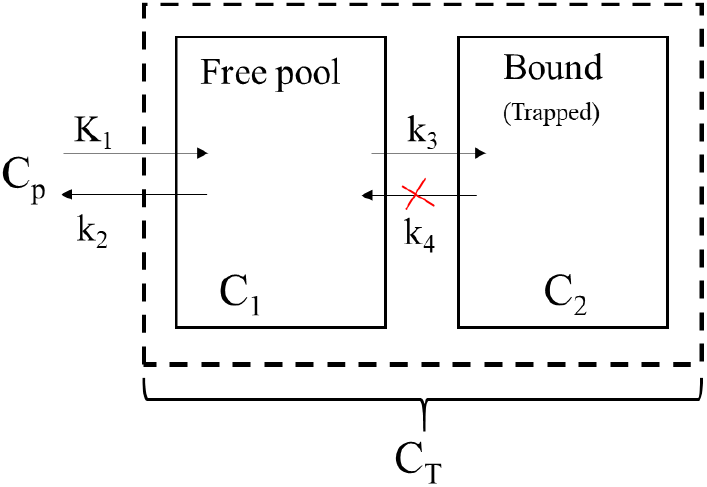
The Two-tissue compartmental model for irreversible PET tracer quantification with two tissue (C_1_, C_2_) and one blood compartment (C_p_) and 3 parameter rate constants (K_1_, k_2_ and k_3_) with the second compartment (C_2_) considered irreversible or trapped (k_4_ = 0)

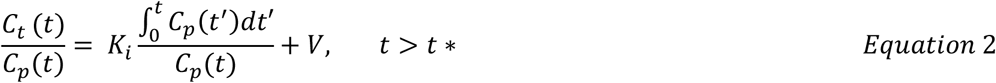

where *K*_*i*_ is the net influx rate of tracer from blood plasma to tissue at each voxel (*K*_1_*k*_3_/(*k*_2_ + *k*_3_)) in min^-1^, *V* is the total blood volume of the tracer initial distribution in the reversible compartment at each voxel, *C*_*p*_*(t)* is the blood plasma tracer concentration at time *t* - the IDIFs, *C*_*t*_*(t)* is the measured tissue timed activity curve at each voxel, *t* is the mid-time points of each frame of the dynamic PET scan, and *t*^*\**^ is the time after tracer injection when the reversible compartment is relatively in steady state with blood plasma. At steady state, *K*_*i*_ can be determined for a given voxel as the slope of the regression plot of the linear relationship between the ratio of *C*_*t*_*(t)* to *C*_*p*_*(t)* concentrations and *t′* the so-called ‘normalized’ or ‘stretched’ time – ratio of the time integral of the blood plasma concentration to the blood plasma concentration at time t. The tracer distribution volume (DV) can be determined from the intercept of the plot, as the combination of the tracer volume of the reversible compartment and the blood volume. In caliPER, the user defines *t*^*\**^ using the control panel parameter entry dialog to set the earliest time when tracer equilibrates with tissue.

For FDG, a whole brain map or regional values describing CMRglc (μmol/100g/min) can be generated by multiplying *Ki* by the individual measured pre-injection blood glucose (mmol/L) normalized by the lump constant (0.52) (Reivich et al., 1985).

## 3. Evaluation and Application

### 3.1 Image Acquisition

Retrospective PET/MRI from 19 healthy volunteers and 8 frontotemporal dementia patients from 2 studies acquired between April 2014 and July 2017 were used to evaluate the performance of caliPER. All participants provided written informed consent and the study protocol was approved by the Western University Health Sciences Research Ethics Board and conducted in accordance with the Declaration of Helsinki ethical standards. Animal experiments to validate caliPER IDIF with serial arterial sampling were conducted according to the guidelines of the Canadian Council on Animal Care and approved by the Animal Use Committee at Western University. Both animal and human imaging were performed on a clinical PET/MRI scanner (Biograph mMR, Siemens Healthcare GmbH, Erlangen, Germany) using a 16-channel (12-head; 4-neck array) PET-compatible MRI coil. The head of all participants were immobilized, and a 60 min dynamic PET acquisition in list-mode was performed immediately following FDG injection at 5 MBq/kg. High resolution three-dimensional (3D) anatomical T1-weighted and/or time-of-flight (TOF) arterial angiography MRI scans were acquired in conjunction with PET imaging for IDIFs and spatial normalization for group-level analysis of PET data.

### 3.2 Validation of PVE Correction Approach in A Porcine Model

Nine juvenile female Duroc pigs (18.84 ± 4.82 kg; blood glucose = 3.5 ± 1.9 mmol/L; 70.6 ± 14.1 MBq) were scanned under an anaesthetic combination of isoflurane (1%–3%) and an intravenous infusion of propofol (6–25 mL/kg/h). FDG injection were administered as a slow bolus (15-30 seconds) via a cephalic vein and arterial blood were sampled from the femoral arteries during PET/MR imaging. An MRI compatible automated blood sampling system (Twilite two, Swisstrace GmbH) calibrated to the PET/MRI scanner was used for continuous blood sampling (temporal resolution of 1s) via a withdrawal pump at a sampling rate of 4 ml/min for the first 5 mins, 1ml/min for the next 5 mins, and 0.5 ml/min for remaining 50 mins. The arterial blood data were processed (decay correction and calibration of Twilite to the PET/MRI scanner) and converted to AIFs by rebinning to match the reconstructed PET time frames, using the Psample software (PMOD Technologies Ltd.). The AIFs were then corrected for dispersion and time-shifted using MATLAB (version 2018a, The MathWorks, Natick, MA). The PET data were reconstructed to 51 image volumes (2s × 15 frames; 5s × 6;15s x 8; 30s × 4; 60s × 5; 120s × 5; 300s x 8; matrix size, 344 ×344 ×127; voxel size, 0.8 × 0.8 × 2 mm) using the Siemens e7 tools and an iterative reconstruction algorithm without point-spread function modelling (ordered subset expectation maximization (OSEM) 4 iterations and 21 subsets, zoom factor of 2.5) and smoothed by a 2-mm gaussian filter. The PET data were corrected for decay, scatter, and dead-time, while attenuation correction was performed using computed tomography. The 3D TOF MRI angiography sequence consisted of a matrix size =640 × 640 × 136; voxel size = 0.3 × 0.3 × 0.6 mm; TE =3.6 ms; TR = 21 ms and flip angle = 18°, while the T1-weighted images were acquired using the magnetization prepared rapid gradient echo (MPRAGE) sequence (matrix size = 256 × 256 × 176; voxel size = 1 × 1 × 1 mm; TE = 2.98 ms; TR = 2000 ms; flip angle = 9°). The IDIFs were extracted from the internal carotid arteries in the neck using caliPER as described above including PVE corrections. The corrected IDIFs generated from TOF and MPRAGE vessel segmentation were compared to dispersion-corrected and time-shifted AIFs to evaluate the level of performance of using the staple T1-weighted imaging as anatomical reference compared to TOF imaging that are not standard in brain imaging but provide enhanced contrast for identifying vessels. The performance of the IDIFs were evaluated by comparing the area under the curve (AUC) generated using Matlab between IFs. Bias was estimated as the ratio of the area under the curve, defined as AUC (IDIF) / AUC (AIF) ∼ 1.

### 3.3 Evaluation of Performance of the PET/MR IDIF

IDIFs were generated from the petrous segment of the internal carotid arteries of eleven healthy younger human volunteers (44 ± 16 years old; 5 females; 204 ± 39.32 MBq; blood glucose = 5.08 ± 0.51 mmol/L) using caliPER (supplementary figure S3). The FDG injection was administered as a bolus injection. The PET data were reconstructed to 51 image volumes (2s × 15 frames; 5s × 6;15s x 8; 30s × 4; 60s × 5; 120s × 5; 300s x 8; matrix size, 344 ×344 ×127; voxel size, 0.8 × 0.8 × 2 mm) using the Siemens e7 tools and an iterative reconstruction algorithm without point-spread function modelling (ordered subset expectation maximization (OSEM) with 3 iterations and 21 subsets, zoom factor of 2.5) and smoothed by a 2-mm gaussian filter. The PET data were corrected for decay, scatter, and dead-time. Attenuation correction was performed using the vendor-provided ultrashort echo time MRI sequence and an offline attenuation map generation approach (RESOLUTE) (Ladefoged et al., 2015)). The 3D TOF sequence (voxel size = 0.6 × 0.6 × 0.7 mm; TE = 3.6 ms; TR = 22 ms flip angle = 18°) and T1-weighted MPRAGE sequence (matrix size = 256 × 256 × 240; voxel size = 0.8 × 0.8 × 0.8 mm; TE = 2.25 ms; TR = 2400 ms; flip angle = 8°) were acquired during PET scanning. To validate the PET/MRI IDIFs to the reference blood sampling approach, population-based input functions (PBAIFs) were generated in lieu of individual AIF using AIFs from ten healthy controls described in (Sundar et al., 2018). The FDG blood activity data obtained from Sundar et al (2018) were normalized by the ratio of the individual injected dose and weight (SUV) prior to averaging across the subjects. The PBAIFs were interpolated to match the IDIFs time points and scaled using the cross-calibration factor between the blood sampling system (Twilite two, Swisstrace GmbH) and the PET/MRI scanner. Although our PET scanning protocol and system are similar to Sundar et al, we calibrated the PBAIF individually by scaling the venous blood activity (in SUV) measured in our healthy human controls at 45 and 60 mins post-injection to minimize residual inter-site variations. The individuals IDIFs generated using caliPER were converted from Bq/ml to SUVs. Biases in IDIFs were estimated as described above using the ratio of the area under the curve. To assess whether biases in IDIFs translate to similar biases in quantification of K_i_ and/or CMRglc, global gray matter (GM) and white matter (WM) K_i_ and CMRglc values were calculated for each participant using the IFs and whole brain GM and WM tissue time activity curves in Matlab. The subject-specific GM and WM masks aligned to corresponding mean late frame (45-60 min) PET images were derived from the individual T1-weighted tissue segmentation in SPM 12 (http://www.fil.ion.ucl.ac.uk) with the tissue segmented image thresholded to include voxels with tissue probabilities >0.9 or >0.8, respectively.

### 3.4 Application of caliPER in Frontotemporal Dementia Imaging

To assess the potential utility of caliPER in clinical imaging applications, eight patients with FTD (80 ± 17 years old; 4 females; 190 ± 42.33 MBq; blood glucose = 5.54 ± 1.79 mmol/L) and eight older healthy volunteers (77 ± 16 years old; 4 females; 189 ± 16.55 MBq; blood glucose = 4.75 ± 0.57 mmol/L) recruited from the Cognitive Neurology Clinic at Parkwood hospital (London, ON, Canada) were scanned using the PET/MR imaging protocols described above, except no TOF MRI was acquired. Detailed description of the demographics and clinical profile of this study cohort were previously summarized (Anazodo et al., 2018). PET reconstruction and image processing in caliPER to derive IDIFs using T1-weighted MRI and generate pixel-wise CMRglc parametric maps were performed as described above. To demonstrate the potential added value of absolute tracer quantification using this blood-free approach, the effect size of the regional CMRglc differences between the patient cohort and controls were compared to the clinical standard relative SUV approach. SUV images were generated from 30 to 45-minute post-injection scans and reconstructed to one image volume using parameters described above. The SUV and CMRglc images were registered to the corresponding T1-weighted anatomical image, skull stripped, spatially transformed to the Montreal Neurological Institute (MNI) standard space using the unified segmentation-based normalization approach in SPM (Ashburner and Friston, 2005), and spatially smoothed using 8 mm FWHM gaussian kernel. The spatially smoothed SUV images in standard space were count normalized by the mean SUV value in the occipital lobe to minimize known inter-subject variabilities. Regions-of interest analysis was performed on the CMRglc and relative SUV (rSUV) images in key brain regions associated with FTD (Anazodo et al., 2018).

### 3.5 Statistical Analysis

To test for differences in performance of the two MR techniques for PET/MR IDIF, the AUC ratio for the pig model and healthy human controls were compared using two sample t-tests. Because the PVE and spillin corrections are dependent on how well the vessels are delineated and on the user’s ROI selection (figure 1), an inter-observer agreement analysis was performed between two users at two institutions to test for observer-dependent factors that could influence the PET/MR IDIF performance. Concordance in the mean vessel diameters for IDIF in the pig and younger healthy controls were assessed between users using the Interclass correlation coefficient (ICC). ICC estimates and their 95% confidence intervals were calculated based on a mean-rating (*k*=2), absolute-agreement, and a two-way random effects model. ICC values >0.90 indicate excellent reliability/agreement. The global K_i_ and CMRglc values for the PBAIFs and the two IDIFs were compared in younger healthy controls using a one-way ANOVA, individually for GM and WM. Pearson correlations and Bland-Altman plots were performed to further probe for biases in quantification using IDIFs compared to PBAIFs. The mean CMRglc and rSUV in key brain regions were compared between FTD patients and controls using a two-sample t-tests. The measured effect sizes of the changes observed in CMRglc and rSUV between-group comparisons were computed using Cohen’s d effect size test. All statistical analysis was performed in SPSS (IBM, Armonk, NY, United States, version 27) and statistical significance was set for statistic values with p<0.05.

## 4. Results

caliPER was tested on Linux (Ubuntu 16.04) and Microsoft Windows 10 operating systems. The processing time from uploading PET and MRI to generation of parametric maps (K_i_ and CMRglc) take ∼10 min on a typical desktop computer (2.5 GHz Intel core i5/8GB DDR4 RAM/64-bit Linux), depending on the choice of anatomical MRI data. Using TOF images, parametric maps can be created in <10 mins because of high vessel-to-soft tissue contrast. The automated processing steps were successful completed on all dataset used for evaluation and validation, without any failed attempts.

The average internal carotid artery diameter measured from pigs was 4.44 ± 0.31 mm, yielding an average RC for PVE correction of 0.21 ± 0.01 and average spillin correction factor of 0.67 ± 0.01. Figure 4 highlights representative PVE corrected IDIF curves and the cumulative AUCs in agreement with the corresponding AIF in the porcine model and PBAIF in a young healthy control. The ICC (2,2) = 0.954 (0.790, 0.991) between the two users showed no systematic difference in measured pig vessel sizes between users. Overall, both TOF and MPRAGE based IDIFs were overestimated by 8% (TOF = 8 ± 5%; MPRAGE = 8 ± 14%) and performed equally with no statistically significant differences observed in the AUC ratios (figure 4c), demonstrating that even in relatively small arterial vessels, IDIFs can be effectively extracted using our approach including PVE corrections. In the human young volunteers the measured internal carotid diameters in the petrous segment were on average 6.11 ± 0.66 mm, corresponding to a mean RC of 0.31 ± 0.04 and a mean spillin coefficient of 0.57 ± 0.03 (supplementary figure S4, table S1). The corrected IDIFs were in general within the range of PBAIFs as demonstrated in figure 4d for one individual. The ICC (2,2) = 0. 975 (0.898, 0.994) between the two users showed excellent agreement and reliability in measured vessel sizes between users. On average the TOF IDIFs overestimated the IFs by 6 ± 8% while the MPRAGE IDIFs were overestimated by 6 ± 9.8% (figure 4f), as observed in the ratio of the AUCs between the IDIFs and PBAIFs.

**Figure 4:**
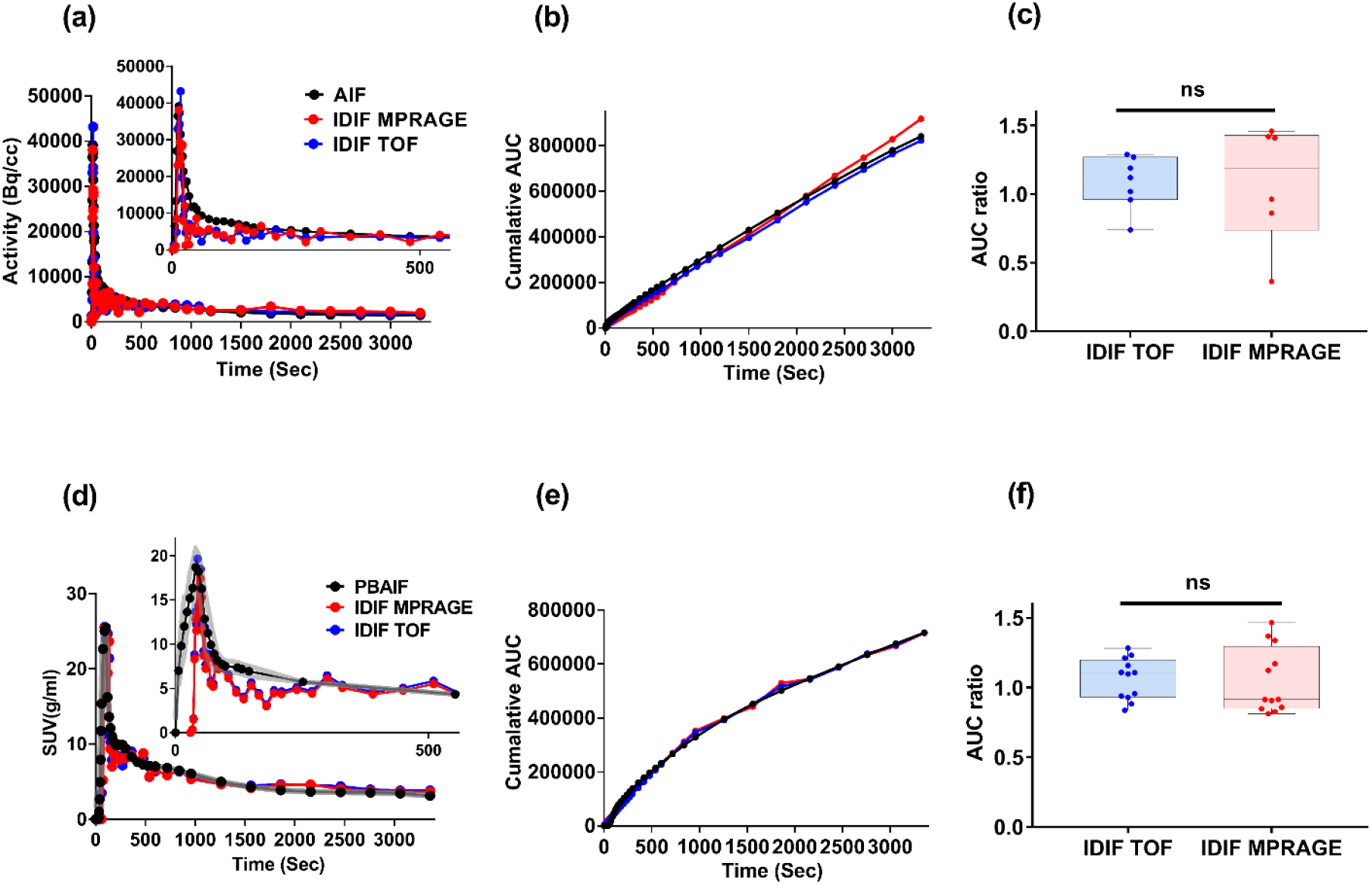
PVE and spillin corrected PET/MR IDIFs generated using TOF (blue curves) and MPRAGE MRI (red curves) produce similar IFs to blood-derived data (black curves) in the pig model (**top row**) and in younger human controls (**bottom row**). Example of IFs in an individual pig and human volunteer (**a, d**) demonstrate good agreement between IDIFs and the AIF or PBAIF as shown in the individual’s corresponding cumulative AUC (**b, e**). The IDIFs for the human controls were in general within range of the PBAIFs (gray curves). Overall, the AUC ratios across participants (pigs N=9; younger humans N=11) indicate similar performance between the two MR IDIFs with no significant (ns) difference (**c, f**).

Although the IDIFs appear to overestimate the measured IFs, there were no observed statistically significant differences in global mean whole brain K_i_ and CMRglc values in pig model as well as in the global mean GM and WM K_i_ and CMRglc values in the younger healthy controls when IDIFs were used in the Patlak graphical analysis compared to AIFs or PBAIFs (figure 5, supplementary figure S5). Across the 11 younger healthy participants, global mean (median) GM and WM CMRglc were 24.72 ± 5.40 (23.7) μmols/100g/min and 14.7 ± 3.6 (14.02) μmols/100g/min for TOF IDIF, 24.60 ± 4.85 (24.15) μmols/100g/min and 15.40 ± 2.70 (15.35) μmols/100g/min for MPRAGE IDIF and 26.17 ± 4.90 (25.7) μmols/100g/min and 14.21 ± 3.5 (13.9) μmols/100g/min for PBAIFs, respectively. Pearson correlation analysis revealed a strong correlation between IFs in global mean K_i_ GM (r = 0.90, p = 0.0001) and WM (r = 0.77, p = 0.004) and GM and WM CMRglc estimates (GM: r = 0.94, p = 0.0001; WM: r=0.84, p = 0.01) (figure 5c, e). The Bland-Altman assessment indicated minimal bias between global GM and WM CMRglc using IDIFs (figure 5d, f), suggesting that IDIFs can provide similar regional brain quantification measures as invasive arterial blood sampling approaches (PBAIF).

**Figure 5.**
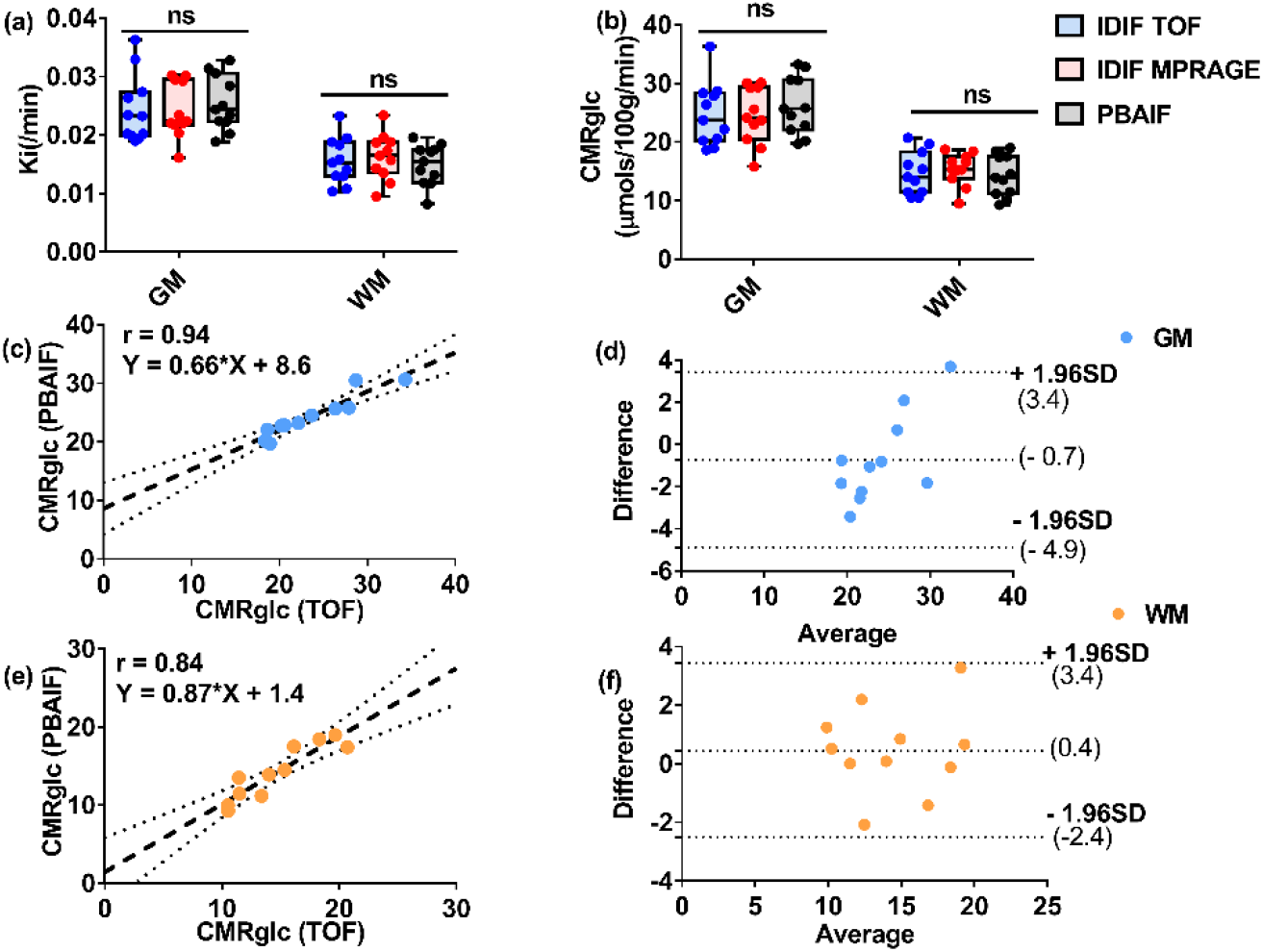
Global gray matter (GM) and white matter (WM) parameters describing the net flux rate, K_i_ (**a**) and cerebral glucose metabolic rate, CMRglc (**b**) in younger healthy controls (N=11) are not significantly (ns) different when IFs are derived from PET/MR IDIFs or PBAIFs. A strong agreement between IDIF and PBAIF GM and WM CMRglc estimates can be seen in the regression plots (p < 0.001) (c, e), while the Bland-Altman plots (d, f) reveal minimal biases.

CMRglc was significantly reduced (p<0.05) across all regions-of-interests in FTD patients compared to controls as shown in figure 6b and illustrated in one patient and control (figure 6 a). Most notably, we observed a significant reduction of CMRglc in the cerebellum and occipital lobes of these patients. In contrast, the relative SUV values measured within the same brain regions were significantly reduced in global GM (t = 2.66, p = 0.03), right insula (t = 4.26, p = 0.004 value), right inferior frontal gyrus (t = 3.57, p = 0.009), and cerebellum (t = 3.47, p = 0.011) in patient cohort (figure 6c). Larger Cohen’s d effect sizes (*d* >0.8) were observed in CMRglc measures in all regions-of-interest (supplementary table S2) while for SUVs the effect sizes were in the larger range for the right insula, right superior temporal gyrus, and cerebellum measurements, but medium in the rest of brain regions including globally in GM and WM (supplementary table S2).

**Figure 6.**
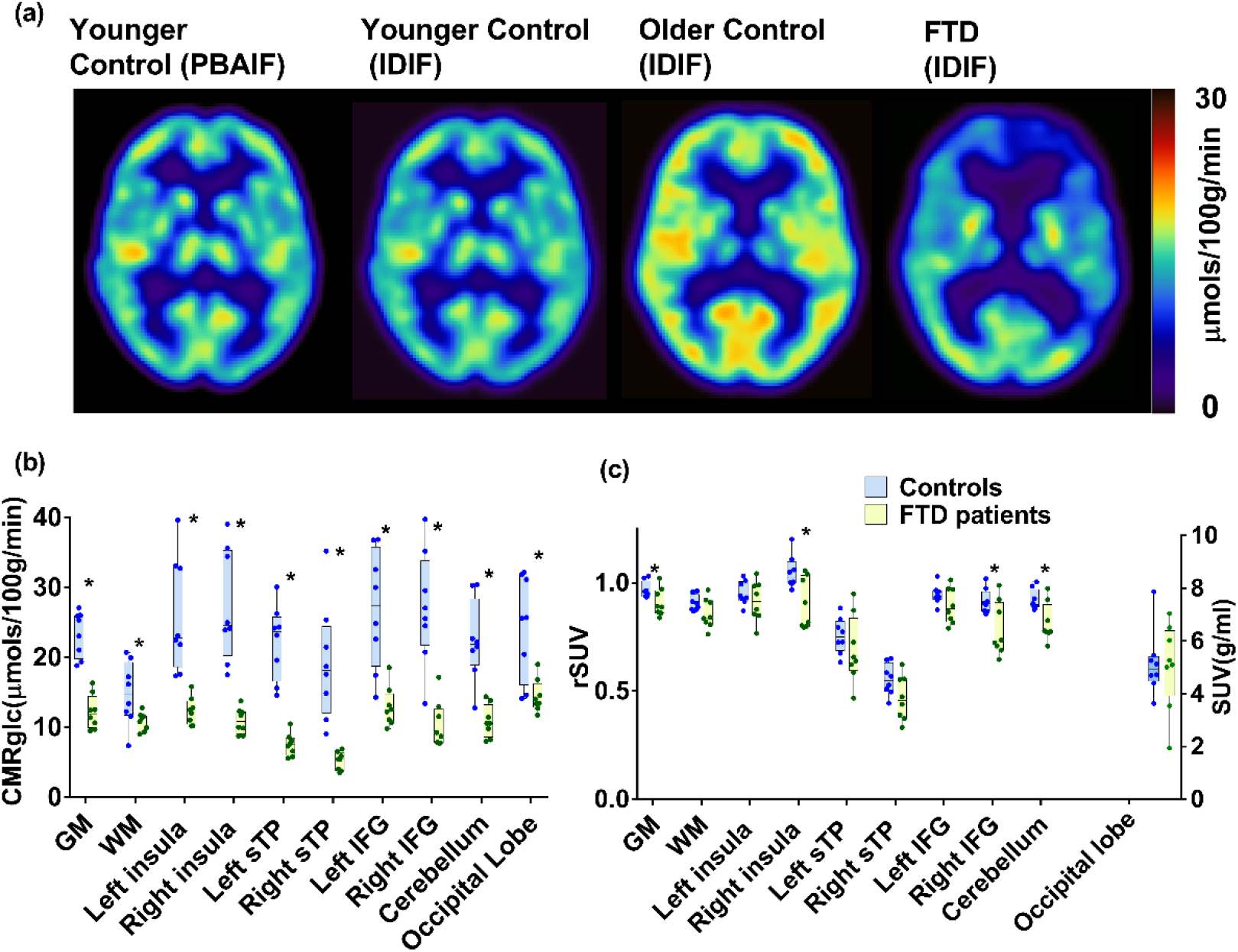
Representative age-and sex-matched CMRglc maps (**a**) generated using caliPER and T1-weighted IDIF in a patient with FTD (58-year-old, female), an individual from the older healthy control group (59-year-old, female) and an individual from the younger healthy control group (60-year-old, female). For comparison, the younger healthy control maps compared visually to maps generated using an established PET modelling tool (PMOD) and PBAIF. Significantly reduced CMRglc (**b**) are observed bilaterally in key FTD brain regions; insula, superior temporal gyrus (sTP) and inferior frontal gyrus (IFG) as well as in the cerebellum (p<0.05). In contrast, FDG uptake measured as SUVs (**c**) were significantly reduced globally in gray matter (GM) and some regions (*). SUVs were normalized by the occipital lobe, which was not statistically different between groups.

## 5. Discussion

PET is considered the gold standard imaging modality for measuring physiologic functions and neurochemistry because of its ability to accurately estimate parameters describing the uptake and distribution of tracers in tissue. While PET quantification fundamentally provides meaningful biological information (e.g. regional glucose consumption in μmol/100g/min) and can effectively characterize target tissue function independent of other physiological factors (e.g. tissue perfusion or target receptor availability), PET imaging is routinely performed using semi-quantitative methods such as SUVs that provide no meaningful biological information, in order to eliminate the invasive arterial blood sampling required for absolute PET quantification. This trade-off between precision (direct parametric mapping) and feasibility (semi-quantification), degrades the intrinsic value of PET and limits its use as the fundamental standard mode for in vivo molecular imaging. We addressed this inherent trade-off by implementing and evaluating a PET/MR IDIF approach called caliPER, a high-fidelity prototype image processing software for generating relevant parametric maps (K_i_, DV) to enable absolute PET quantification of clinical imaging.

caliPER is intended as a clinical image processing tool and as such was developed on the MeVisLab rapid prototyping platform, which features an extensive leading-edge library of algorithms for image segmentation, registration, volumetry analysis, and multi-dimensional visualization. The modular framework within MeVisLab provides integration of new algorithms/modules scripted in Python for advanced image processing and analysis such as our implementation of the Patlak graphical analysis, as well as development of simplified image processing workflows in a user-friendly graphical user interface, familiar in clinical environments. All the required libraries and packages to install and run caliPER are pre-complied such that it can be easily installed as a standalone software in Linux, Microsoft Windows, or macOS by simply running the executable file (.exe). Unlike other available PET image processing and quantification tools, caliPER is a comprehensive software that takes reconstructed dynamic PET DICOM images and generates parametric pixel-wise maps without the need for externally pre-processed input functions, blood-/image-derived (c.f. supplementary table 3). All the necessary image processing steps to generate subject-specific parametric maps are provided within caliPER including options for manual segmentation and registration when automated options fail. Although caliPER was optimized for PET/MR IDIF using simultaneously acquired PET and MRI data, its 2-step image co-registration approach makes it feasible to also use MRI acquired separately from different session/scanner. This co-registration approach also permits the use of any subject-specific anatomical MRI with sufficient tissue contrast to discriminate vessels as we previously demonstrated for body imaging in [^68^Ga]-PSMA-11 imaging using a T1-weighted volumetric interpolated breath-hold examination (VIBE) 3-point Dixon sequence (water image) where vessels appear bright enough at sufficient resolutions even on the 18-second Dixon MR attenuation scans (Ringheim et al., 2020), and in [^18^F]-NaF imaging using a T2-weighted half-Fourier-acquired single-shot turbo spin echo (HASTE) sequence where blood vessels appear dark (Cui *et al* 2019), as well as here in this study using TOF and MPRAGE sequences.

caliPER is currently designed to generate parametric maps of net influx rate for tracers with irreversible uptake or binding, and specifically for FDG, the pixel-wise measure of cerebral metabolic rate of glucose consumption. By using a porcine model with an internal carotid vessel size that are on average just below the spatial resolution limit of the Siemens mMR system (4.3 mm FWHM) (Delso et al., 2011), we demonstrated that the simple empirically determined PVE corrections implemented in caliPER can effectively recover blurred PET signal and produce IDIFs with relatively minimal bias compared to the gold standard arterial blood sampling. However, PVE correction factors may need to be derived for other PET isotopes because of the variable positron contribution, which may degrade the measured PET activity due to scattering events. For very short-lived isotopes such as ^15^O, correction factors derived using ^13^N or ^11^C can be substituted given the similar branching ratios. The use of empirically derived PVE correction factors may not be ideal for blood-free IDIF estimation, because the cylindrical phantoms may not robustly simulate the shape of arteries to precisely estimate the correction factors (Zanotti-Fregonara et al., 2011). This could be an important source of bias in producing accurate kinetic parameters in older adults or individuals with vessel disease where vessel shape and diameter are pathologically altered. However, this can be mitigated if 1) high resolution (1mm^3^ isotropic) MRI is used to segment and estimate the measured arterial diameter, and 2) an ideal segment of the vessel where the vessel is straight and cylindrical is carefully selected as the vessel ROI mask, as is implemented in caliPER (supplementary figures S3 and S4 and table S1). It should be noted that underestimating the arterial diameter by 1 mm would decrease the CMRglc estimate by roughly 17% (Croteau et al., 2010). Since PVE correction is necessary to produce IDIF curves approximating AIF curves, we chose this simpler approach instead of other approaches such as region-based methods (Sari et al., 2016; Sundar et al., 2018) to account for both spill-out and spill-in effects. Region-based approaches derive a recovery coefficient to account for spill-in effects ignoring spill-out effects and do so by solving the complex geometric transfer matrix of measured and ‘true’ (corrected) PET activity, based on the assumption of uniform activity within each region. The definition of regions that contribute to PVE in each voxel highly influence the accuracy of the derived recovery coefficient. Much care is required when defining regions for region-based approaches to ensure that all potential sources of spill-in activity to the vessel region are included during the computation of correction factors. Poor estimation of spill-in effects can lead to an overestimation of the vessel radioactivity concentration and bias IDIFs, particularly at the tail-end of the curve. For IDIFs which are best suited for graphical-based kinetic analysis, overestimating the tail-end of the curve further minimizes the accuracy of parametric mapping.

Overall, the IDIF extracted using caliPER approximated the AIFs obtained from direct arterial blood sampling, with a small error. The K_i_ and CMRglc estimated using IDIF were similar to estimates using blood sampling and within limits of whole brain values reported by others, considering differences in lump constant for CMRglc (c.f. supplementary table S4) (Chen *et al* 1998, Croteau *et al* 2010, Reivich *et al* 1985, Sundar *et al* 2018, Zanotti-Fregonara *et al* 2009). The absolute median % difference between global gray matter CMRglc values derived from PBAIF and IDIF in the younger healthy volunteers were slightly higher (∼7%) compared to the PET/MR IDIF approach (4%) described by (Sundar *et al* 2018), but well within the 14% difference between repeated AIF measurements in their test-restest study (Sundar *et al* 2018). The performance difference between the two PET/MR IDIF approaches could be a result of the use of PBAIF as an alternate but imperfect reference standard in our retrospective analysis, compared to the prospective AIF analysis by Sundar et al. In lieu of measured AIFs, normalized and calibrated PBAIFs can produce reliable IFs for quantitative FDG imaging in healthy individuals and may serve as an adequate alternative to continuous arterial blood sampling, although with relatively small errors (<5%) (Shiozaki et al., 2000; Takikawa et al., 1993). However, PBAIF may not be adequate for clinical imaging as its performance in patient populations could be impacted by disease pathophysiology, limiting the application of PBAIF derived from healthy individuals or from a specific patient cohort to another group of patients (Zanotti-Fregonara et al., 2013). Nonetheless, there is a general understanding that IDIFs are imprecise, with poor estimation of the arterial peak and a better estimation of the tail-end of the curve (Zanotti-Fregonara et al., 2011), largely due to the inaccuracies in recovery coefficients and sampling of total administered bolus (Chen *et al* 1998). Slower tracer bolus infusion can improve reconstruction of the IFs and minimize possible detector dead-time in capturing the peak count rate. Although rapid bolus (<15s) injections were used in this study for FDG imaging, an intravenous injection infused slowly over 30-40s may better model the shape of the IF. Regardless, the area under the peak is negligible compared to the total area under the curve. More importantly, graphical approaches such as Patlak rely largely on estimation of the steady-state tracer concentration in the tail-end as well as the total area under the curve but not the precise estimation of the shape of the peak (Zanotti-Fregonara *et al* 2011). For reversible tracers such as FDG, underestimation of the peak by up to 20% would translate to an error of <0.1% in CMRglc estimated using Patlak and AIF/IDIF (Chen *et al* 1998).

CMRglc quantification using caliPER demonstrated a larger effect size in detecting differences between the patient and control groups compared to clinical-standard SUVs. A study by (Yamaji et al., 2000) showed equally inferior performance of SUV to CMRglc in detecting disease patterns in Alzheimer’s disease. The authors observed significant CMRglc reductions in several brain regions in patients with mild and moderate Alzheimer’s disease compared to healthy volunteers but found no difference in SUVs between patients with mild Alzheimer’s disease and controls and no SUV changes in the moderate Alzheimer’s disease group in some brain regions detected by CMRglc (Yamaji et al., 2000). In oncology, Patlak or compartmental modelling parameters have been shown to give smaller (more focal) tumour volume measures (Visser et al., 2008) and provide better predictors of treatment response and outcome (Dunnwald et al., 2011). Given the limited number of studies with direct comparison of SUV and Patlak quantification parameters, especially at subject-specific level, the propensity for improved detection of disease patterns using IDIF and Patlak needs to be systematically investigated before blood-free quantification approaches can be used as suitable alternative to clinical-standard SUVs.

Our approach for IDIF can be further improved by compensating for motion artefacts in the dynamic PET data. Patient motion degrades PET image quality and can artificially bias the measured activity, producing inaccurate parameter estimates in PET tracer kinetic models (Zanotti-Fregonara et al., 2012). Perhaps this is one reason the IDIF generated using caliPER has slightly larger bias compared to the PET/MR IDIF method recently reported by (Sundar *et al* 2018) where head motion in PET data was compenstated. Several approaches have been introduced for motion correction of PET including the use of external motion tracking devices, or MRI-based motion-tracking techniques to estimate motion and apply correction retrospectively after PET data acquisition (Catana, 2015). Retrospective motion correction approaches can effectively correct patient motion, if motion vectors can be optimally estimated throughout the lengthy PET acquisition. We previously demonstrated the feasibility of using ultra-fast MRI pulses embedded as navigator echoes in anatomical MRI sequences to retrospectively correct motion in brain PET and anatomical MRI, including relatively large head movements of up to 11° rotations and 14 mm translations (Johnson et al., 2019). MRI navigators can be incorporated into serial anatomical and functional MRI acquisitions during standard PET/MR imaging to capture motion throughout the dynamic PET scan without increasing MRI scan time. As more approaches for motion correction become widely available including emerging machine learning techniques, motion-corrected PET and MRI can be imported into caliPER.

As next steps, we plan to use the modular framework of MeVisLab to extend caliPER functionalities to include 1) conversion of whole blood IDIFs to plasma IDIFs using population-based blood-to-plasma data, 2) implement simplified kinetic analysis for reversible tracers (e.g. TSPO PET) using suitable graphical analysis methods (Zhou et al., 2009), 3) fully automated PET/MRI segmentation and registration processing steps to minimize potential user-operator dependent bias, and 4) automated regions-of-interest analysis for brain imaging using a brain atlas. As currently implemented, regions-of-interest analysis can be performed by manually drawing ROIs on the pixel-wise parametric maps.

## 6. Conclusion

In general, caliPER can produce subject-specific blood-free image-derived input functions similar to input functions generated with complete blood sampling. The ability to derive parametric maps using Patlak graphical analysis for simple dynamic kinetic modelling shows promise with respect to clinical feasibility. Unlike available kinetic modelling software, caliPER provides an all-in-one tool for non-invasive kinetic modelling, from reconstructed PET data to parametric maps without the need of deriving IDIFs elsewhere. By creating a comprehensive user-friendly simple tool for blood-free absolute PET quantification, we provide a means to increase widespread adoption of kinetic analysis for absolute PET quantitation. Further validation of caliPER including the IDIF approach using other PET tracers and on different scanners and settings (sequential vs. simultaneous MRI acquisition) is still required to establish the utility and value of caliPER for kinetic analysis of PET data.

## Acknowledgements

This study was funded by research grants from the Healthy Brains, Healthy Life (HBHL; 2bNISU17), Weston Foundation (RR182074), Canadian Institutes of Health Research (148600), and Mitcas Accelerate Fellowship (IT06382). The authors would like to thank Linshan Liu, John Butler, Heather Biernaski, and Laura Morrison for assistance with experimental data collection. The authors acknowledge the contribution of Pei-Shan Wei in implementing the input function fitting routine. The authors are grateful to Lalith Sundar and Thomas Beyer for sharing their arterial input functions data for population-based input function analysis.

## Data And Code Availability

In this work, caliPER was used as a standalone version. This research tool and its functionalities as presented here can be accessed through the *syngo*.via Frontier platform (version VB40). caliPER will be uploaded to the Siemens Healthineers Digital Marketplace and made available to users after creating and registering a free account (https://store.teamplay.siemens.com/apps). It will also be made available to the Open Applications Digital Ecosystem, allowing potentially for improvement in speed of processing and for closer integration within clinical workflows (i.e., hybrid 3D viewing, neurological volumetric analysis, and oncology image analysis and reading). Access to the caliPER standalone version along with sample data can be provided by an email request to.

## Author Contribution – CRediT Roles

Conceptualization and study design: UA, CL, and GM.

Data Collection and Curation: UA, MK, KsL, EF, and AS.

Software development and visualization: UA, CL, SZ, and GM.

Software validation and data analysis: PD, CL, and UA.

Supervision and Project Administration: UA

Original manuscript draft: PD, CL, and UA.

Writing-review and editing: All authors and co-authors

Funding acquisition: KsL and UA

