## Supplementary Materials for "caliPER: A software for blood-free parametri*c* P*a*t*l*ak mapp*i*ng using *PE*T/M*R*I input function"

**Figure S1**: A screen capture of caliPER's Front-end illustrating its use in body imaging data analysis. Here in Patlak analysis of [$^{68}$Ga]-PSMA-11.

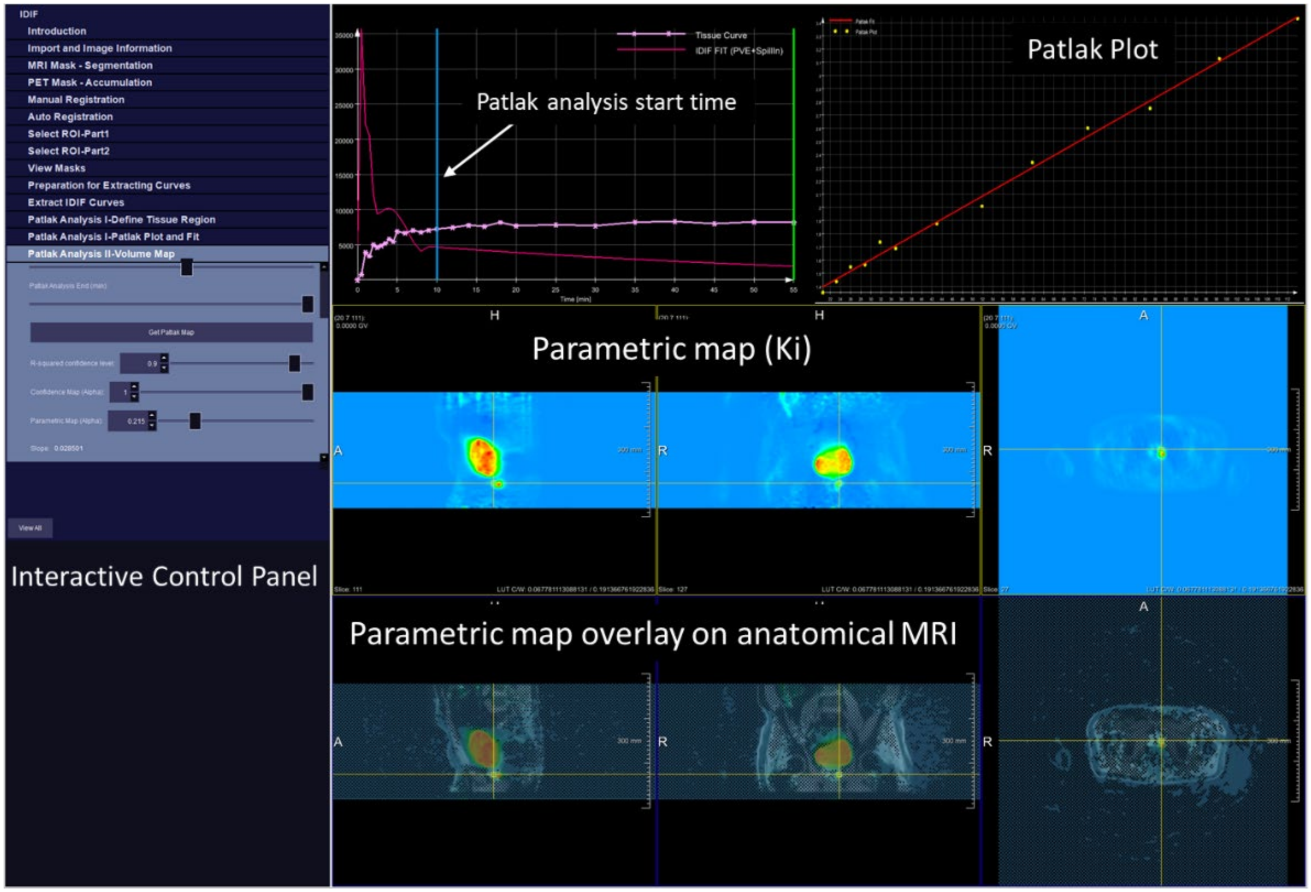

Simulation Experiments for PVE and Spill over Correction Factors

To generate an IDIF corrected for PVE and spillover contamination of non-arterial voxels, a phantom experiment was performed on a PET/MRI scanner (Biograph mMR, Siemens Healthcare, Erlangen, Germany), as described in (Croteau *et al* 2010) and (Feng *et al* 2012). Correction factors as a function of arterial diameter were obtained using the NEMA IEC Body Phantom Set™ containing a hollow torso that simulates background concentration in human body and refitted with six 2.5 cm long cylinders (on ~4 cm posts) with inner diameters ranging from 5 to 30 mm -(figure S1) designed to mimic the arterial geometry and capture the resolution limit of the internal carotid arteries. Although the NEMA Phantom was specifically designed to measure image quality of PET/CT systems, we adapted the phantom for PET/MR imaging by reconstructing PET images from the PET/MRI with an aligned CT attenuation correction map of the phantom acquired on a PET/CT (Discovery VCT, GE Healthcare, Waukesha, USA). This ensured that the image quality and contrast recovery coefficients from PET/MRI, described below and used as correction factors for PVE and spillover contamination were void of bias related to inadequate PET attenuation correction (Ziegler *et al* 2015).

For PVE correction, the phantom was filled with $^{18}$F-FDG at a concentration of ~4:1 sphere-to-background activity ratio, where all spheres were uniformly aliquoted 46.6 kBq/ml calibrated to the start of the PET scan (a concentration within limits of what is typically administered to subjects in clinical studies) (Croteau *et al* 2010), while the cooler background in the torso had 12.1 kBq/ml. The phantom was scanned for ten minutes during acquisition of a T2 turbo spin echo image (0.7 x 0.7 x 4.0 mm), acquired to help delineate the contours of the cylinders. The phantom experiment was repeated within a week, using the same radioactivity concentration. To simulate spillover contamination, the phantom was scanned as described above using cold cylinders and 25 kBq/ml of FDG in the torso compartment to mimic the spillover of radioactivity from non-arterial voxels into the lumen of vessels. All images were reconstructed to one volume using similar reconstruction parameters for the validation and evaluation studies described below; 3D ordinary Poisson ordered-subset-expectation-maximization with 3 iterations, 21 subsets, 344 matrix size, and 2 mm Gaussian post-smoothing filter using the Siemens e7 tools.

To derive the recovery coefficients (RC) as a function of vessel diameter for PVE correction, the maximum PET activity in the hottest slice was extracted from circular regions of interest (ROI) placed around each cylinder, carefully delineated using T2-MRI ROI masks and then fitted to the model in equation S1 (Croteau *et al* 2010, Feng *et al* 2012).

$$RC\ (x) = \frac{a(x-d)}{(1+\left(\frac{x-d}{b}\right)^{c})^{1/c}} \qquad Equation\ S1$$

Where *x* is the diameter of the ROI in mm, *d* is the lowest measured diameter of the ROI in mm and *a, b* and *c* are coefficients.

The resulting RC curve (figure S1) yields correction factors for vessels of known diameter on PET images and can be extrapolated to correct PVE bias from vessels diameters below the simulated cylinder limit (<5 mm). Likewise, the spillover correction factor (figure S1) as a function of vessel diameter was obtained from the mean activity in each cylinder and fitted to model in equation S2 (Feng *et al* 2012).

$$Spillover\ (x) = (1 - Plateau) * \ e^{-(k*w)} + Plateau \qquad Equation\ S2$$

Where *x* is the diameter of the ROI in mm, k is a constant equal to the reciprocal of the spillover range, and the Plateau is the asymptotic spillover contribution.

All phantom image analysis and curve fitting were performed using MATLAB (Version R2018A; MathWorks Natick Massachusetts). The RC and spillover coefficients are time-independent constants ranging from 0 to 1.

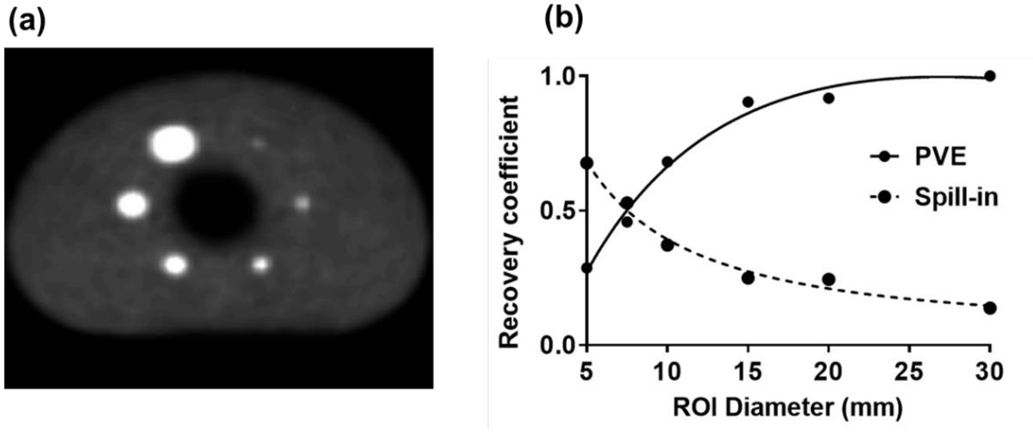


**Figure S2**. (a) A PET image of the refitted NEMA Phantom showing an axial slice through center of the cylinder rods. (b) PVE recovery coefficient (RC) (R-square = 0.99 RMSE = 0.02) and spillin (R-square = 0.97 RMSE = 0.04) correction curves.

**Figure S3. Illustration of the internal carotid vessel tree extracted in caliPER and corresponding IDIF ROIs in two human volunteers.**

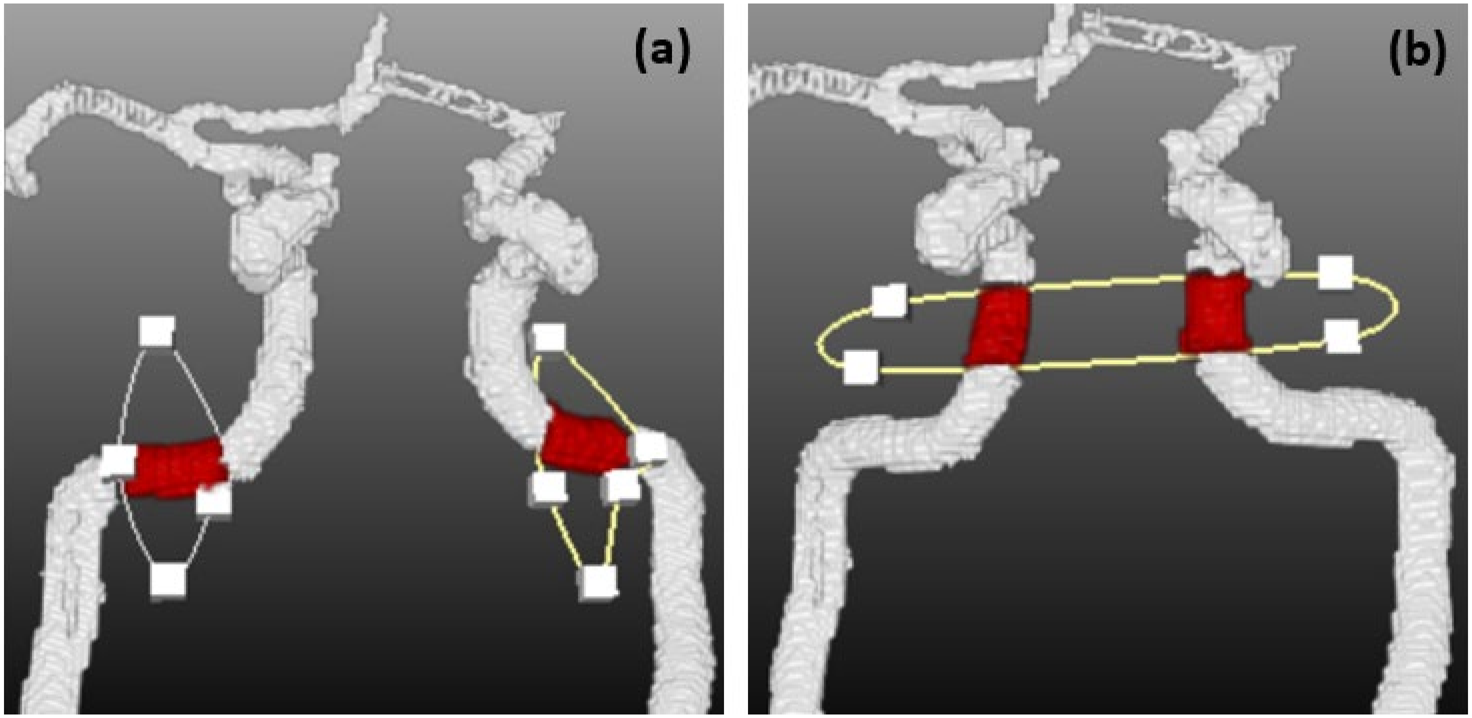


Representative snapshots from the caliPER viewer of the internal carotid arteries in a younger healthy control extracted using TOF MRI depicting the IDIF ROI selected from the petrous segment as single ROI selection (**a**) or alternatively as both arteries selected from the cavernous segment (**b**).

**Figure S4. MRI measures of the internal carotid artery (petrous segment) diameters**

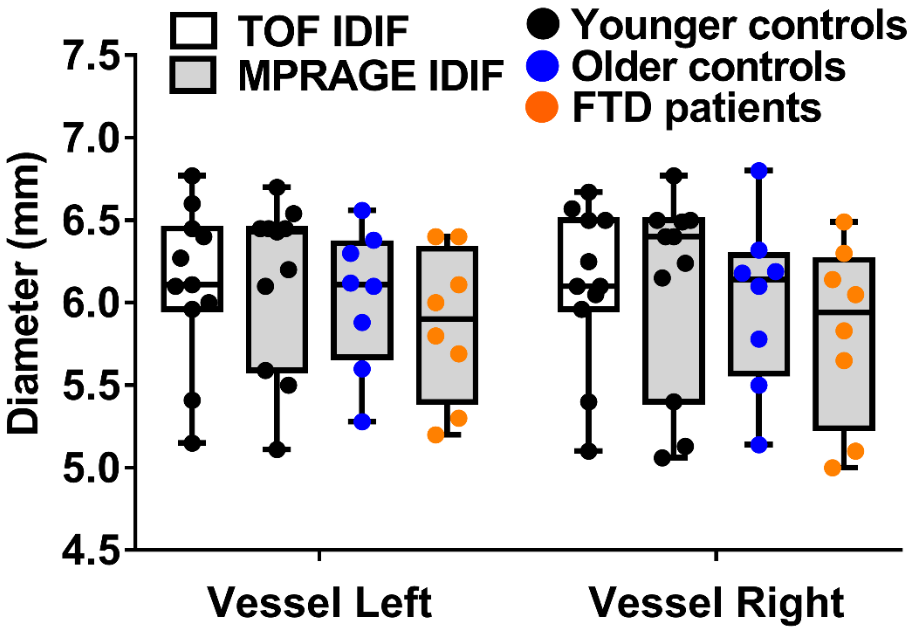


Vessel diameters measured from the left and right petrous segment of the internal carotid artery using TOF and MPRAGE MRI in the younger and older control groups and frontotemporal dementia (FTD) patient group are shown. The average diameter of both carotid arteries measured on MPRAGE MRI in the older control (6.01 ± 0.46 mm) and FTD patient (5.84 ± 0.48 mm) groups relative to the younger controls (6.11 ± 0.54 mm) were slightly smaller (table S4 below) but not statistically significantly different between groups. Compared to the average diameter in the younger control group, the smaller vessel size in the FTD patient group was estimated to produce a 3.92% bias in Ki/CMRglc.

### Figure S5: Global mean whole brain (WB) net flux rate (Ki) and cerebral glucose metabolic rate in the pig model

The global mean WB $K_i$ derived from TOF IDIF, MPRAGE IDIF and AIF were 0.016 ± 0.003 $min^{-1}$, 0.015 ± 0.007 $min^{-1}$ and 0.015 ± 0.005 $min^{-1}$, respectively. The global mean WB CMRglc derived from TOF IDIF, MPRAGE IDIF and AIF were 18.35 ± 4.75 µmol/100g/min, 17.8 ± 8.94 µmol/100g/min, and 17.64 ± 7.35 µmol/100g/min, respectively. The relative % difference in Ki and CMRglc derived using both IDIFs were 6.67% and 4.30%, respectively and no statistical difference between input functions were observed, as shown in figure S4 below.

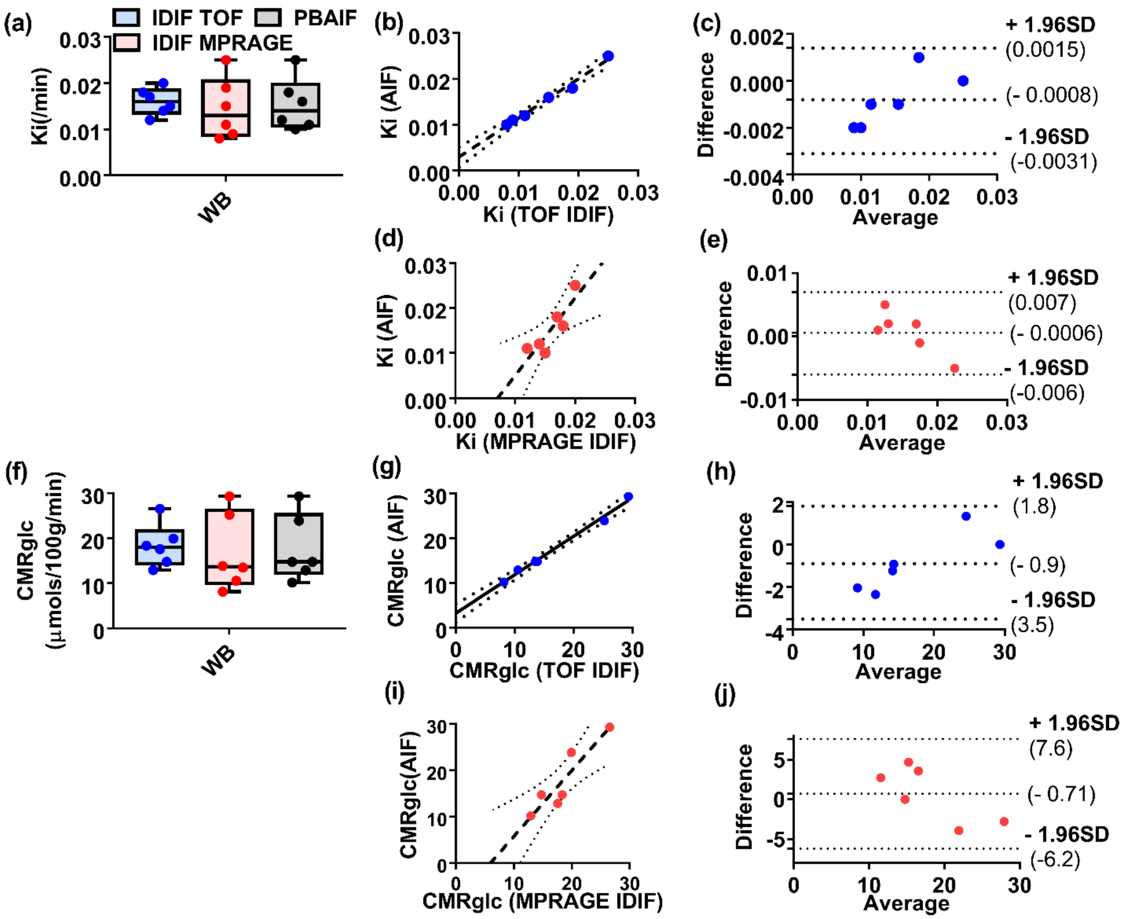


Figure S5: Global mean whole brain (WB) parameters describing the net flux rate, $K_i$ (**a-e**) and cerebral glucose metabolic rate, CMRglc (**f-j**) in the pig model. The parameters generated from the IDIFs (MPRAGE and TOF) were in good agreement with parameters generated with the AIFs and no statistically significant difference in mean values were observed ($p>0.05$). The whole brain $K_i$ estimates showed strong agreement between input functions (TOF (**b**): $r^2 = 0.99$, slope = 0.85, intercept = -0.003; MPRAGE (**d**): $r^2 = 0.77$, slope = 1.71, intercept = -0.01) and minimal bias (**c and e**). Similarly, the whole brain CMRglc measures showed strong agreement between input functions (TOF (**g**): $r^2 = 0.99$, slope = 0.86, intercept = 3.2; MPRAGE (**i**): $r^2 = 0.84$, slope = 1.42, intercept = -8.5) and minimal bias (**h and j**).

**Table S1: Mean and standard deviation vessel diameter, PVE and spill-in correction factors**

| Cohort | Younger Controls (n=11) | Older Controls (n=8) | FTD Patients (n=8) |
|---|---|---|---|
| Left Carotid (mm) | 6.14 ± 0.51 | 6.03 ± 0.42 | 5.90 ± 0.50 |
| Right Carotid (mm) | 6.10 ± 0.60 | 6.00 ± 0.51 | 5.80 ± 0.54 |
| PVE correction factor | 0.31 ± 0.05 | 0.30 ± 0.06 | 0.30 ± 0.04 |
| Spill-in | 0.58 ± 0.04 | 0.58 ± 0.04 | 0.59 ± 0.03 |

*Measurements in T1-MPRAGE are shown. PVE and RC correction factors from average of the left and right vessel diameters are shown. For completeness, the mean vessel diameter in the younger control group measured using TOF-MRI for left and right carotid were 6.11 ± 0.48 and 6.11 ± 0.49, respectively.*

**Table S2: Effect size of group differences between patients with FTD and Healthy age-matched controls**

| Brain Region | Effect Size (Cohen's d) | |
|---|---|---|
| | CMRglc | rSUVs |
| **Gray matter** | 3.57 | 0.71 |
| **White matter** | 1.22 | 0.89 |
| **Left insula** | 2.48 | 0.50 |
| **Right insula** | 3.37 | 1.47 |
| **Left superior temporal gyrus** | 1.69 | 0.52 |
| **Right superior temporal gyrus** | 2.32 | 0.96 |
| **Left Inferior frontal gyrus** | 1.54 | 0.79 |
| **Right Inferior frontal gyrus** | 2.57 | 1.33 |
| **Cerebellum** | 5.13 | 1.47 |

### Table S3: List of available packages and scripts for PET tracer kinetic modelling

| *Feature* | *Open source* | *Operating System* | *Language* | *Quantification* | *Default IF* | *Processing* | *Visualization* | *Reference* |
|---|---|---|---|---|---|---|---|---|
| ***Qmodelling*** | Yes (SPM) | Mac, Linux, Windows | MATLAB | SRTM, Patlak, Logan | Reference Tissue | Local | GUI | López-González *et al* 2019[1] |
| ***APPIAN*** | Yes (Github) | Linux | NiPype | SRTM, Patlak, Logan | AIF | Web | Dashboard | Funck *et al* 2018[2] |
| ***PPET*** | Yes | Linux, Windows | IDL VM | SRTM, Patlak, Logan, Spectral analysis | AIF | Web | GUI | Boellaard *et al* 2006 |
| ***Multiparametric PET Suite AI*** | No | Siemens syngo | | Patlak for FDG | IDIF, AIF, PBIF | Local | syngo | Siemens Healthcare GmbH[3] |
| ***Voxulus*** | No | Philips | MATLAB | SRTM, 1-&2-TCM | AIF, IDIF | Local | IntelliSpace | Philips GmBH [4] |
| ***Magia*** | Yes (Github) | Mac, Linux | MATLAB | SRTM, Patlak, Logan, 1-&2-TCM | AIF/Reference Tissue | Local | GUI | Karjalainen *et al* 2020[5] |
| ***DEPICT*** | Yes | Linux, Windows | MATLAB | 1-&2-TCM | AIF | Local | GUI | Gunn *et al* 2002[6] |
| ***KiWi*** | Yes (Github) | Linux, Mac, Windows | MATLAB | 1-TCM | AIF | Local | GUI | Phan *et al* 2017 [7] |
| ***COMKAT*** | Yes | Mac, Linux | MATLAB | 1-&2-TCM | AIF | Local | GUI | Muzic *et al* 2001[8] |
| ***MIAKAT*** | Yes | Mac, Linux | MATLAB | 1-&2-TCM, SRTM | AIF | Local | GUI | Gunn *et al* 2016 |
| ***Vienna IDIF*** | Yes | Linux, Windows | MATLAB | Patlak | IDIF*[a] | Local | - | Sundar *et al* 2019 |
| ***TKMF*** | Yes | Mac, Linux | MATLAB | 1-/2-TCM, Patlak, | AIF | Web | - | Huang *et al* 2005[9] |

| | | | | | | | | |
|---|---|---|---|---|---|---|---|---|
| | | | | Logan | | | | |
| ***kinfitr*** | Yes (Github) | Linux, Windows, Mac | R | 1-&2-TCM, Patlak, Logan | AIF | Local | - | Matheson 2019[10] |
| ***PetSurfer*** | Yes | Linux, Mac | Tcl/Tk | MRTM1, MRTM2, Logan | Reference Tissue | Local | - | Greve et al 2014 |
| ***PMOD*** | No | Mac, Linux, Windows | JAVA | 1-&2-TCM, Patlak, Logan | AIF | Local | GUI | PMOD Technologies LLC[11] |
| ***caliPER*** | Siemens Frontier | Mac, Linux, Windows | C++ | Patlak | IDIF*[b] | Local | GUI | |

[a]Ability to import high MRI for IDIF (time of flight (TOF) validated); [b]Ability to import MRI for IDIF (validated on several anatomical MRI including TOF and T1-weighted sequences); IF: input function; SRTM: simplified reference tissue model; AIF: arterial input functions; IDIF: image derived input functions. *Corrected for partial volume errors including spill-in errors; IDL VM: IDL virtual machine, 1-TCM: 1-tissue compartment model; 1-&2-TCM: 1- or 2- tissue compartment model; MTRM: Multilinear Reference Tissue Model

Note tools such as MIKAT, Magia, Appian, and PMOD can used externally derived IDIF for modelling, while IDIF can be derived within PMOD simply by drawing regions of interest on PET data, but without partial volume errors correction. Multiparametric PET Suite AI herein are not commercially available in all countries. Its future availability cannot be guaranteed. Please contact your local Siemens Healthineers organization for further details

[1] http://www.uimcimes.es/contenidos/golink?p=1

[2] https://github.com/APPIAN-PET

[3] https://www.siemens-healthineers.com/molecular-imaging/options-and-upgrades/software-applications/flowmotion-multiparametricpet-suite

[4] https://www.philips.com/a-w/imalytics/specialized-modules/pharmacokinetic-modeling

[5] https://github.com/tkkarjal/magia

[6] http://www.bic.mni.mcgill.ca/~rgunn/DEPICT.html

[7] https://github.com/jennyannphan/KiWi

[8] http://comkat.case.edu/index.php?title=Home

[9] https://dragon.nuc.ucla.edu/modelfitting/ (*was not accessible at the time of manuscript submission*)

[10] https://github.com/mathesong/kinfitr

[11] https://www.pmod.com/web/

### Table S4: Reported net influx rate ($K_i$) and glucose consumption rate (CMRglc) values in healthy volunteers from the Literature

| **Study** | **n/age/M** | **Glucose concentration (mmol/L)** | **Lump constant (LC)** | **Influx constant (Ki; min$^{-1}$)** | **CMRglc reported** | **CMRglc (µmol/100g/min)**[§] | **CMRglc (Adjusted to LC = 0.52) (µmol/100g/min)** |
|---|---|---|---|---|---|---|---|
| **Stender., *et al* (2015)** | 29/ 44 ±16 /19 | 4.97±0.56 (89.5 ± 10.1 mg/dL) | 0.61 | | PBAIF-WB: 0.30 ± 0.031 µmol/g/min<br>PBAIF-BS: 0.23 ± 0.03 µmol/g/min | PBAIF-WB: 30 ± 3.1<br>PBAIF-BS: 23 ± 3.0 | PBAIF-WB: 35 ± 3.6<br>PBAIF-BS: 26.9 ± 3.5 |
| **Croteau., *et al* (2010)** | 6 / 59 ±23 /5 | | 0.52 | AIF: 0.029 ± 0.011<br>IDIF: 0.039 ± 0.007 | AIF-FL: 13.8 ± 3.9 mg/100g/min<br>IDIF-FL: 14.0 ± 5.7 mg/100g/min | AIF-FL: 76.6 ± 21.6<br>IDIF-FL: 77.0 ± 31.35 | AIF-FL: 76.6 ± 21.6<br>IDIF-FL: 77.0 ± 31.35 |
| **Sari., *et al* (2017)** | 18 / 28.3 (22-40)/ 18 | 5.07 (4.3-5.7) | 0.89 | AIF: 0.049 ± 0.004<br>IDIF: 0.046 ± 0.008 | AIF-GM: 26.88 ± 2.36 mg/100g/min<br>IDIF-GM: 25.38 ± 4.66 mg/100g/min | AIF-GM: 149.2 ± 13.1<br>IDIF-GM: 139.6 ± 25.63 | AIF-GM:255.4 ± 22.4<br>IDIF-GM: 238.9 ± 43.9 |
| **Sundar., *et al* (2019; ~~2020~~)** | 10 / 27 ± 7 /5 | 4.61 -6.32 | 0.65 | | AIF-WB: 32.0 ± 6.0 µmol/100g/min<br>IDIF-WB: 32.0 ± 6.0 µmol/100g/min<br>AIF-SF: 34.4 ± 6.60 | AIF-WB: 32.0 ± 6.0<br>IDIF-WB: 32.0 ± 6.0<br>AIF-SF: 34.4 ± 6.60 | AIF-WB: 40.0 ± 7.5<br>IDIF-WB: 40.0 ± 7.5<br>AIF-SF: 43±8.25 |
| **Hahn**., *et al* (2016)** | 15 / 24 ± 4 /7 | | 0.89 | | AIF-PC: 22.7 ± 4.4 µmol/100g/min<br>AIF-HP: 19.4 ± 3.1 µmol/100g/min | AIF-PC: 22.7 ± 4.4<br>AIF-HP: 19.4 ± 3.1 | AIF-PC: 38.8 ± 7.5<br>AIF-HP: 33.1 ± 5.3 |
| **Huisman *et al*., (2012)** | 9/ 36 ± 13 / 9 | 5.5 ± 0.2 | 0.52 | AIF GM: 0.031 ± 0.004<br>AIF WM: 0.010 ± 0.0008 | AIF-GM: 29 ± 3 µmol/100g/min<br>AIF-WM: 19 ± 1 µmol/100g/min | AIF-GM: 29 ± 3<br>AIF-WM: 19 ± 1 | AIF-GM: 29 ± 3<br>AIF-WM: 19 ± 1 |

| | | | | | | | |
|---|---|---|---|---|---|---|---|
| **Roelcke *et al* (1997)** | 16/40 ± 15/7 | | | | AIF-WB: 43.6 ± 6.9 µmol/100g/min | AIF-WB: 43.6 ± 6.9 | AIF-WB: 43.6 ± 6.9 |
| **Huang et al (1980)** | 13/-/- | 5.1 ±0.53 (91.9 ± 9.6 mg/dL) | 0.42 ± 0.06 | AIF-GM: 0.33 ± 0.006 | AIF-GM: 7.3 ± 1.18 mg/100g/min | AIF-GM: 40.2 ± 6.5 | AIF-GM: 32.5 ± 5.2 |
| | | | | AIF-WM: 0.015± 0.004 | AIF-WM: 3.41 ± 0.64 mg/100g/min | AIF-WM: 18.8 ± 3.5 | AIF-WM: 15.2 ± 2.8 |
| **Reivich et al (1985)** | Rate constant: 9 /18-25/ 9 | | 0.52 ± 0.03 | AIF-GM: 0.035[ǂ] | AIF-WB: 5.66 ± 0.37 mg/100g/min | AIF-WB: 31.1 ± 2.0 | AIF-WB: 31.1 ± 2.0 |
| | CMRglc: 6 /19-30/ 6 | | | AIF-WM: 0.023[ǂ] | | | |
| **caliPER** | 11/ 44 ± 16/ 5 | 5.08 ± 0.51 | 0.52 | PBIF-GM: 0.025 ± 0.005 | PBIF-GM: 26.2 ± 4.9 | PBIF-GM: 26.2 ± 4.9 | PBIF-GM: 26.2 ± 4.9 |
| | | | | PBIF-WM: 0.015 ± 0.003 | PBIF-WM: 14.21 ± 3.5 | PBIF-WM: 14.21 ± 3.5 | PBIF-WM: 14.21 ± 3.5 |
| | | | | IDIF-GM: 0.024 ± 0.005 | IDIF-GM: 24.6 ± 4.9 | IDIF-GM: 24.6 ± 4.9 | IDIF-GM: 24.6 ± 4.9 |
| | | | | IDIF-WM: 0.016 ± 0.004 | IDIF-WM:15.4 ± 2.7 | IDIF-WM:15.4 ± 2.7 | IDIF-WM:15.4 ± 2.7 |
| | 8/ 77 ± 16 / 4 | 4.75 ± 0.57 | | IDIF-GM: 0.024 ± 0.004 | IDIF-GM: 23.34 ± 3.2 | IDIF-GM: 23.34 ± 3.2 | IDIF-GM: 23.34 ± 3.2 |
| | | | | IDIF-WM: 0.015 ± 0.008 | IDIF-WM: 14.83 ± 4.52 | IDIF-WM: 14.83 ± 4.52 | IDIF-WM: 14.83 ± 4.52 |

*Brain regions: Grey matter (GM), White matter (WM), Frontal (FL), Superior frontal (SF), Whole brain (WB), Precentral (PC), Hippocampus (HP), Cortex (CT) and Brain stem (BS); **Constant-infusion FDG injection. [ǂ]Derived from estimates from Huisman et al 2012 three parameter model. [§]Conversion from 1gram of FDG to 1 molar mass of FDG = 181.1495g/mol. This is not an exhaustive table. Note the average whole-brain CMRglc calculated from four studies performed between 1949 and 1972 using the Kety-Schmidt technique is 5.35 ± 0.33 mg/100g/min or29.4 ± 1.80 µmol/100g/min, as summarized in Reivich et al (1985). caliPER MPRAGE IDIF parameters values are listed.*

Huisman, Marc C., Larissa W. van Golen, Nikie J. Hoetjes, Henri N. Greuter, Patrick Schober, Richard G. Ijzerman, Michaela Diamant, and Adriaan A. Lammertsma. "Cerebral Blood Flow and Glucose Metabolism in Healthy Volunteers Measured Using a High-Resolution Pet Scanner." *EJNMMI Research* 2, no. 1 (2012/11/20 2012): 63.

Karjalainen, Tomi, Jouni Tuisku, Severi Santavirta, Tatu Kantonen, Marco Bucci, Lauri Tuominen, Jussi Hirvonen*, et al.* "Magia: Robust Automated Image Processing and Kinetic Modeling Toolbox for Pet Neuroinformatics." [In English]. *Frontiers in Neuroinformatics* 14, no. 3 (2020).

López-González, Francisco J., José Paredes-Pacheco, Karl Thurnhofer-Hemsi, Carlos Rossi, Manuel Enciso, Daniel Toro-Flores, Belén Murcia-Casas, Antonio L. Gutiérrez-Cardo, and Núria Roé-Vellvé. "Qmodeling: A Multiplatform, Easy-to-Use and Open-Source Toolbox for Pet Kinetic Analysis." *Neuroinformatics* 17, no. 1 (2019): 103-14.

Matheson, G. J. Kinfitr: Reproducible PET Pharmacokinetic Modelling in R. *bioRxiv: 755751* (2019). https://doi.org/10.1101/755751

Muzic, R. F., Jr., and S. Cornelius. "Comkat: Compartment Model Kinetic Analysis Tool." [In eng]. *J Nucl Med* 42, no. 4 (Apr 2001): 636-45. Matheson, Granville J. "Kinfitr: Reproducible Pet Pharmacokinetic Modelling in R." *bioRxiv* (2019): 755751.

Phan, JA., Landau, A.M., Jakobsen, S. *et al.* Radioligand binding analysis of $\alpha_2$ adrenoceptors with [$^{11}$C]yohimbine in brain in vivo: Extended Inhibition Plot correction for plasma protein binding. *Sci Rep* **7,** 15979 (2017). https://doi.org/10.1038/s41598-017-16020-1

Reivich, M., A. Alavi, A. Wolf, J. Fowler, J. Russell, C. Arnett, R. R. MacGregor*, et al.* "Glucose Metabolic Rate Kinetic Model Parameter Determination in Humans: The Lumped Constants and Rate Constants for [18f]Fluorodeoxyglucose and [11c]Deoxyglucose." *Journal of Cerebral Blood Flow & Metabolism* 5, no. 2 (1985): 179-92.

Roelcke, U., L. Kappos, J. Lechner-Scott, H. Brunnschweiler, S. Huber, W. Ammann, A. Plohmann*, et al.* "Reduced Glucose Metabolism in the Frontal Cortex and Basal Ganglia of Multiple Sclerosis Patients with Fatigue: A 18f-Fluorodeoxyglucose Positron Emission Tomography Study." [In eng]. *Neurology* 48, no. 6 (Jun 1997): 1566-71.

Sari, H., K. Erlandsson, I. Law, H. B. W. Larsson, S. Ourselin, S. Arridge, D. Atkinson, and B. F. Hutton. "Estimation of an Image Derived Input Function with Mr-Defined Carotid Arteries in Fdg-Pet Human Studies Using a Novel Partial Volume Correction Method." [In English]. *Journal of Cerebral Blood Flow and Metabolism* 37, no. 4 (Apr 2017): 1398-409.

Stender, Johan, Ron Kupers, Anders Rodell, Aurore Thibaut, Camille Chatelle, Marie-Aurélie Bruno, Michael Gejl*, et al.* "Quantitative Rates of Brain Glucose Metabolism Distinguish Minimally Conscious from Vegetative State Patients." *Journal of Cerebral Blood Flow & Metabolism* 35, no. 1 (2015): 58-65.

Sundar, L. K. S., O. Muzik, L. Rischka, A. Hahn, I. Rausch, R. Lanzenberger, M. Hienert*, et al.* "Towards Quantitative 18f Fdg-Pet/Mri of the Brain: Automated Mr-Driven Calculation of an Image-Derived Input Function for the Non-Invasive Determination of Cerebral Glucose Metabolic Rates." [In English]. *Journal of Cerebral Blood Flow and Metabolism* 39, no. 8 (2019): 1516-30.

Ziegler, S., B. W. Jakoby, H. Braun, D. H. Paulus, and H. H. Quick. "Nema Image Quality Phantom Measurements and Attenuation Correction in Integrated Pet/Mr Hybrid Imaging." [In eng]. *EJNMMI Phys* 2, no. 1 (2015): 18.
